# Modeling Oncolytic Vaccinia Virus Therapy Highlights Neutrophil Impact on Tumor Suppression ^∗^

**DOI:** 10.1101/2024.11.06.622313

**Authors:** Sahaj Satani, Hana Dobrovolny

## Abstract

Oncolytic vaccinia viruses (OVVs) present a promising approach for melanoma treatment due to their ability to selectively infect and lyse tumor cells. In this study, we use an ordinary differential equation (ODE) model of tumor growth inhibited by OVV activity to parameterize previous research on the effect of neutrophil depletion in B16-F10 melanoma tumors in mice. We find that the data are best fit by a model that accounts for neutrophil-mediated viral clearance, and that neutrophil depletion provides a mechanism for enhanced OVV efficacy and tumor reduction. We also find that parameter estimates for the most effective OVV regime share characteristics, most notably a low viral clearance rate by neutrophils, that might explain the improved outcomes. Further studies examining the impact of neutrophil modulation across different tumor models may help elucidate the extent to which these findings generalize and inform the design of novel OVV-based cancer therapies.

## 1 Introduction

Cancer remains one of the leading causes of death worldwide, with melanoma accounting for approximately 7,990 deaths and 97,610 new cases projected in the United States for 2024 [Siegel et al.(2024)]. While the 5-year survival rate for early-stage melanoma exceeds 95%, this rate drops dramatically to only 29.8% for stage IV metastatic melanoma [Cronin et al.(2023)]. Traditional treatments such as chemotherapy, radiation, and surgery often result in significant side effects and may have limited efficacy against metastatic disease [Domingues et al.(2018)]. These limitations have driven the search for more targeted and effective therapeutic approaches.

Oncolytic virotherapy has emerged as a promising treatment strategy, with vaccinia virus showing particular potential due to its natural tumor tropism and extensive safety record in human populations [MacKie et al.(2001)]. The FDA’s approval of T-VEC, a modified herpes virus, for treating inoperable melanomas in 2015 has paved the way for other oncolytic virus platforms [Andtbacka et al.(2015)]. However, current oncolytic viruses face significant challenges in clinical settings, with only 20-30% of patients showing durable responses to treatment [Ribas et al.(2017)].

A major obstacle to successful oncolytic virus therapy is premature viral clearance by the innate immune system, particularly by neutrophils, which can comprise up to 70% of circulating immune cells [Summers et al.(2010)]. Neutrophils can rapidly respond to viral infections, with studies showing that they can infiltrate infected tissues within hours and reduce viral titers by up to 90% through phagocytosis and other clearance mechanisms [Drescher and Bai(2013)]. While this response is crucial for defending against pathogenic infections, it can significantly impair the therapeutic efficacy of oncolytic viruses, with studies showing that high neutrophil infiltration correlates with reduced treatment success rates [Chen and Mellman(2017)].

Mathematical models have been instrumental in understanding virus dynamics, with varying levels of complexity incorporated to describe different aspects of virus-host interactions [Wodarz(2001), Komarova and Wodarz(2010), Friedman et al.(2006)]. Simple models often assume perfect virus spread within tumors, while more complex models consider immune responses and spatial heterogeneity [Kim et al.(2015), Bajzer et al.(2008)]. Recent studies have shown that incorporating immune components into these models can better predict treatment outcomes, with some models suggesting that immune-mediated viral clearance can reduce effective virus particle numbers by up to 99% within 48 hours of administration [Zhou et al.(2024), Paiva et al.(2009)].

In this paper, we develop a mathematical model to analyze the impact of neutrophil-mediated clearance on oncolytic vaccinia virus therapy. By fitting our model to previously published data, we quantify the effects of neutrophil depletion on viral efficacy in melanoma treatment. Our work builds upon existing mathematical frameworks while specifically addressing the role of neutrophils in viral clearance, aiming to identify strategies for optimizing oncolytic virotherapy.

## 2 Materials and Methods

### 2.1 Experimental Data

Tumor data were obtained from the paper *Transient inhibition of neutrophil functions enhances the antitumor effect of intravenously delivered oncolytic vaccinia virus* by Zhou et al. [Zhou et al.(2024)], which examined the role of neutrophils in oncolytic vaccinia virus therapy for melanoma. The researchers investigated how neutrophils can reduce the effectiveness of vaccinia virus by phagocytizing it in the bloodstream. To study this, they used multiple experimental groups: For data from Figure 4F, mice were treated with either PBS (control), oncolytic vaccinia virus (VV) alone, anti-LY6G antibody alone (to deplete neutrophils), or a combination of anti-LY6G antibody and VV (LY6G+VV). The anti-LY6G antibody was administered intraperitoneally at 200 g/day/mouse for two consecutive days prior to virus treatment. Virus was then administered intravenously at 1×10 pfu through the tail vein every other day.

For data from Figure 5J, mice were treated with either CAL-101 (a PI3Kinase delta inhibitor) plus PBS, VV alone, or a combination of CAL-101 and VV (CAL101+VV). CAL-101 was administered orally at 10 mg/kg prior to virus injection. As in the previous experiment, virus was delivered intravenously at 1×10 pfu through the tail vein.

Each treatment group consisted of 6-7 mice that had B16-F10 melanoma cells (1×10 cells) implanted subcutaneously in their right flank. Tumor measurements were taken using calipers and volumes were calculated based on the tumor’s dimensions. Mice were monitored until the tumor reached 1000 mm^3^ or showed signs of ulceration, at which point they were euthanized according to protocol. The data represent the average tumor volumes from each treatment group over time. A more detailed explanation of methods and data collection is available in Zhou et al. (2024).

Data from Figures 4F and 5J, which show the tumor growth curves under different treatment conditions, were extracted using WebPlotDigitizer (accessed on 13 June 2024).

### 2.2 Model Fitting and Statistical Analysis

We evaluate two different mathematical models in fitting the treatment data to see if the added complexity of accounting for neutrophils is justified. In the paper, the researchers note that neutrophils can phagocytize the vaccinia virus, making it less effective at killing the tumor. We investigate a model of oncolytic virus therapy that includes a neutrophil immune response:

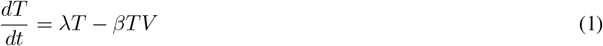

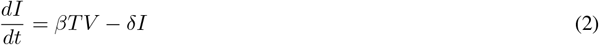

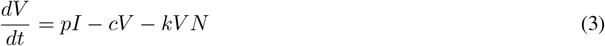

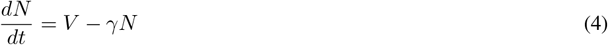

The tumor (*T*) grows at a rate *λ*. The virus (*V*) infects the tumor at rate *β*. The infected cells (*I*) die at a rate *δ* and produce more viral particles at rate *p*. The virus is cleared at baseline rate *c* and through neutrophil-mediated clearance at rate *k*. Neutrophils (*N*) are stimulated by the virus and decay at rate *γ*.

We investigate two conditions: virus alone (VV) and virus with neutrophil depletion through anti-LY6G antibody treatment (LY6G+VV). The researchers reduce the number of circulating neutrophils by treating with anti-LY6G Ab and find that the vaccinia virus is more effective in killing the tumor cells.

In order to estimate the model parameters, we fit the model to experimental data. Model fitting is conducted by systematically varying parameter values to match the model curves to the experimental data. We quantitatively assess how well the model matches the data by calculating the sum of squared residuals (SSR) between the model and observed values.

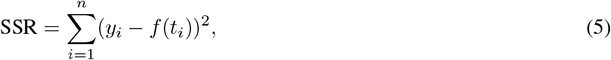

where *n* is the number of data points, *y*_*i*_ is the experimentally observed tumor volume at time *t*, and *f* (*t*) is the model’s predicted tumor mass at time *t*. The parameter values that lead to the minimal value of SSR are considered the best fit parameters estimates.

Since not all viral particles from the initial injection of virus manage to infect cells, we consider *V*_0_, or the initial amount of virus in the system, as a free parameter. Because infection begins when the virus is first introduced to the tumor, we assume the initial number of infected cells *I*(0) and amount of neutrophils *N* (0) are 0. Fitting was conducted in two stages with the control tumor used to estimate the base growth rate. Treated tumors were then assumed to have that same growth rate and other viral parameters were estimated using fits to treated data. The list of parameters that were estimated using this fitting procedure and the biological meaning of the parameters is included in Table 1.

**Table 1:** Model parameters estimated by the fitting process.

| Parameter | Meaning |
| --- | --- |
| $\lambda$ | Tumor growth rate |
| $\beta$ | Viral infection rate |
| $\delta$ | Death rate of infected cells |
| $p$ | Viral production rate |
| $c$ | Baseline viral clearance rate |
| $k$ | Neutrophil-mediated viral clearance rate |
| $\gamma$ | Neutrophil decay rate |

For the fitting of the tumor kinetics model, we use various aspects of the scipy Python package. The ordinary differential model is solved by the odeint function of scipy.integrate, and the minimize function of scipy.optimize minimizes the SSR using the L-BFGS-B algorithm. The virus is administered through multiple injections at days 8, 10, and 12, which is accounted for in the numerical solution by solving the system piecewise between injection times.

We used the L-BFGS-B method from scipy.optimize.minimize to identify the best-fit parameters. Plots were generated using the matplotlib Python package. To determine the 95% confidence intervals for the best-fit parameters and SSR’s, we use 1000 bootstrap replicates. Bootstrapping was also used to determine possible parameter distributions. Distribution graphs were made using pyplot from the matplotlib Python package.

Statistical comparison between VV and LY6G+VV conditions is performed using the Wilcoxon rank-sum test with multiple sampling iterations to ensure robust analysis of parameter differences between conditions. The test is performed on randomly sampled subsets of the bootstrap results to assess the consistency of differences between conditions while accounting for parameter uncertainty.

The model successfully captures the enhanced efficacy of the virus treatment when neutrophils are depleted, supporting the experimental finding that neutrophil-mediated clearance is a significant factor limiting oncolytic virus efficacy. The fitted parameters provide quantitative estimates of the relative importance of different biological processes in determining treatment outcomes.

## 3 Results

### 3.1 Model Fitting Analysis for Standard VV Treatment

We developed a mathematical model incorporating tumor growth, viral infection dynamics, and neutrophil-mediated clearance to analyze oncolytic vaccinia virus therapy. The model was first applied to data from Figure 4F comparing standard vaccinia virus treatment (VV) with neutrophil-depleted treatment using anti-LY6G antibody (LY6G+VV). As shown in Figure 1, our model successfully captured the experimental dynamics for both treatment conditions.

**Figure 1:**
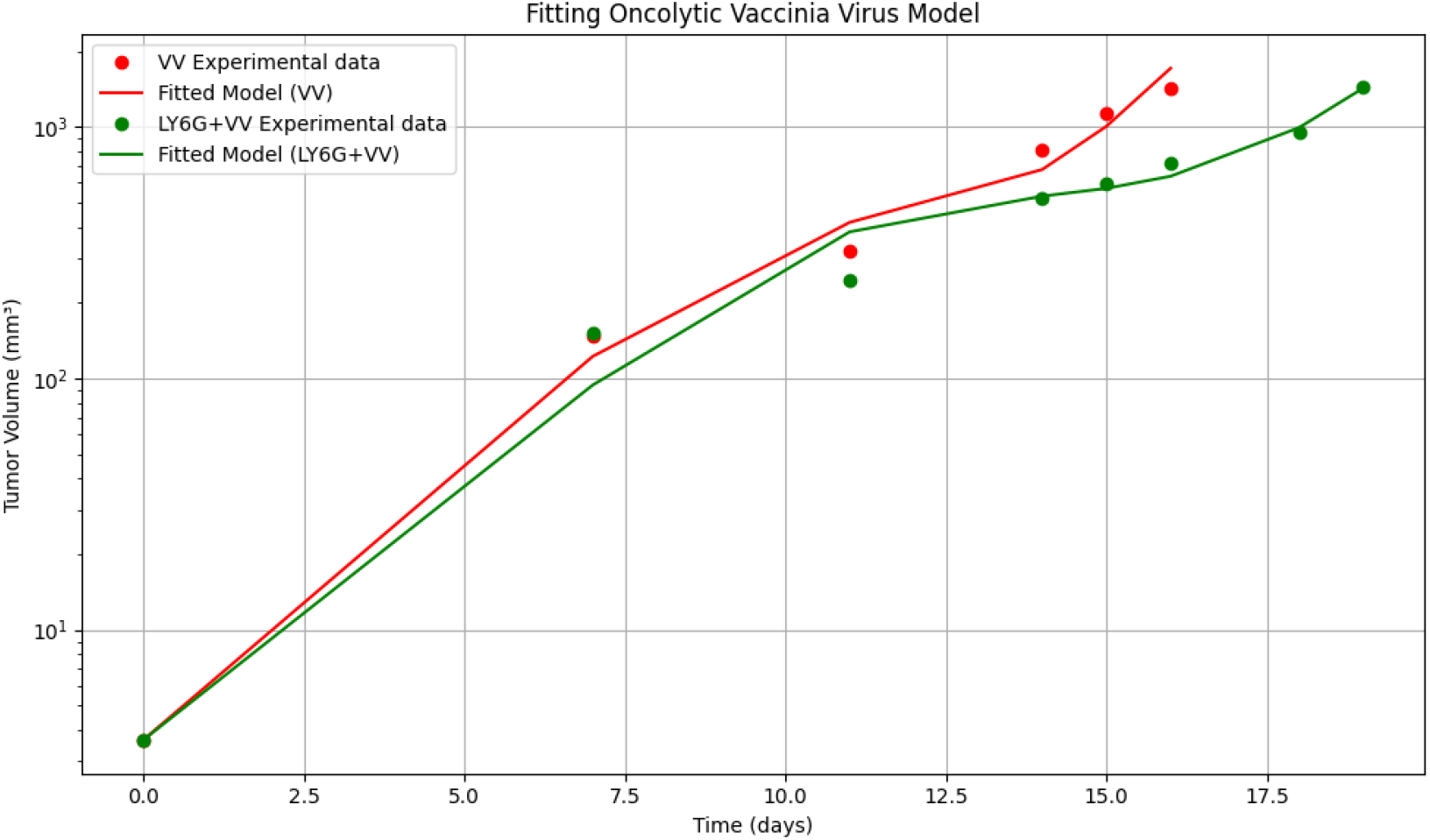
Model fits to experimental data from Figure 4F. Red points and lines show vaccinia virus (VV) experimental data and model fit respectively, while green points and lines show data and model fit for VV treatment with neutrophil depletion (LY6G+VV). The model captures the enhanced tumor suppression with neutrophil depletion. Tumor volumes are plotted on a logarithmic scale.

### 3.2 Parameter Estimates and Bootstrap Analysis

Best-fit parameter values and their 95% confidence intervals revealed substantial differences between treatment conditions (Table 2). Most notably, the neutrophil-mediated viral clearance rate (*k*) differed by four orders of magnitude between conditions (2.29 *×* 10^−6^ for VV vs. 1.45 *×* 10^−10^ for LY6G+VV), quantitatively confirming successful neutrophil depletion. The neutrophil response rate (*γ*) also showed significant variation (2.64 for VV vs. 2.63*×* 10^−2^ for LY6G+VV).

**Table 2:** Best fit parameter values and 95% confidence intervals for the VV and LY6G+VV treatment models from Figure 4F analysis. Parameters represent tumor growth rate (*λ*_*T*_), infection rate (*β*), death rate (*δ*), viral production (*p*), viral clearance (*c*), neutrophil clearance of virus (*k*), and neutrophil clearance rate (*γ*).

| Treatment | Parameter Values | 95% CI | SSR |
| --- | --- | --- | --- |
| VV | $\lambda_T = 6.65 \times 10^{-1}$ | $2.59 \times 10^{-1} - 4.44 \times 10^0$ | 0.0350 |
| | $\beta = 1.13 \times 10^{-7}$ | $1.00 \times 10^{-10} - 1.45 \times 10^{-5}$ | |
| | $\delta = 2.45 \times 10^{-1}$ | $1.00 \times 10^{-3} - 1.00 \times 10^1$ | |
| | $p = 1.01 \times 10^0$ | $3.11 \times 10^{-1} - 2.79 \times 10^0$ | |
| | $c = 1.50 \times 10^{-1}$ | $3.70 \times 10^{-3} - 5.54 \times 10^{-1}$ | |
| | $k = 2.29 \times 10^{-6}$ | $4.03 \times 10^{-9} - 3.05 \times 10^{-3}$ | |
| | $\gamma = 2.64 \times 10^0$ | $9.64 \times 10^{-2} - 1.00 \times 10^1$ | |
| LY6G+VV | $\lambda_T = 7.32 \times 10^{-1}$ | $3.13 \times 10^{-1} - 9.68 \times 10^0$ | 0.0853 |
| | $\beta = 3.70 \times 10^{-9}$ | $6.25 \times 10^{-10} - 9.69 \times 10^{-8}$ | |
| | $\delta = 1.87 \times 10^{-1}$ | $1.00 \times 10^{-3} - 1.00 \times 10^1$ | |
| | $p = 1.00 \times 10^{-1}$ | $6.82 \times 10^{-2} - 1.44 \times 10^{-1}$ | |
| | $c = 1.33 \times 10^{-2}$ | $1.00 \times 10^{-3} - 7.32 \times 10^{-2}$ | |
| | $k = 1.45 \times 10^{-10}$ | $1.00 \times 10^{-10} - 1.43 \times 10^{-8}$ | |
| | $\gamma = 2.63 \times 10^{-2}$ | $1.00 \times 10^{-3} - 1.00 \times 10^1$ | |

Bootstrap analysis of parameter distributions (Figure 2) confirmed these findings, showing clear separation between treatment conditions for neutrophil-related parameters while maintaining similar distributions for tumor growth characteristics.

**Figure 2:**
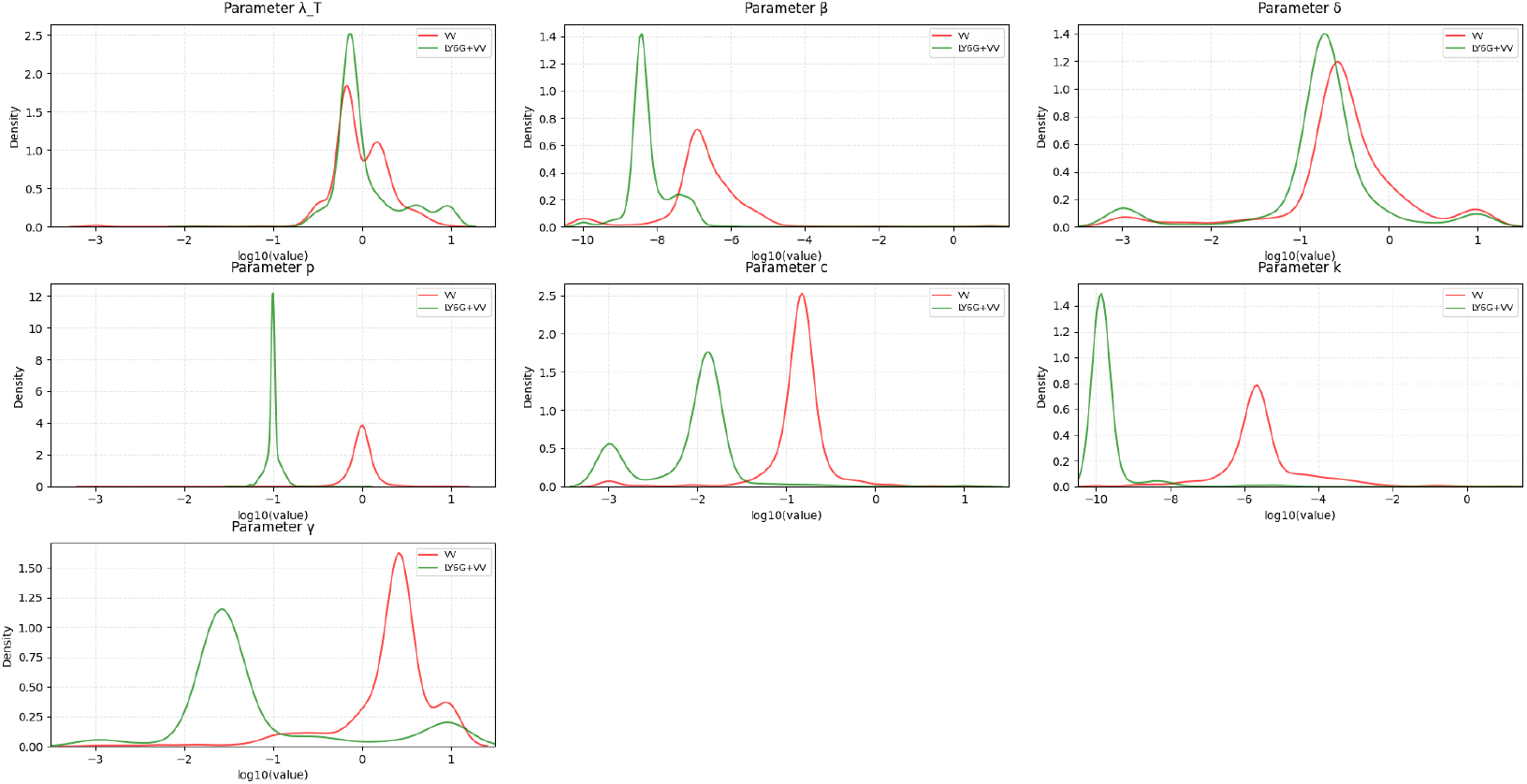
Distribution of bootstrapped parameter values for VV and LY6G+VV treatments from Figure 4F analysis. Parameter distributions show clear separation in neutrophil-related parameters (*k* and *γ*) between treatments. The x-axis represents log10 of parameter values, and y-axis shows density of bootstrap results.

### 3.3 CAL-101 Treatment Effects

To investigate an alternative approach to neutrophil modulation, we applied our model to data from Figure 5J comparing CAL-101 treatment effects. The model successfully captured the dynamics of all three experimental conditions (Figure 3), with SSR values indicating good agreement between model predictions and experimental data (CAL-101+PBS: 0.0672, CAL-101+VV: 0.0463, VV: 0.0329).

**Figure 3:**
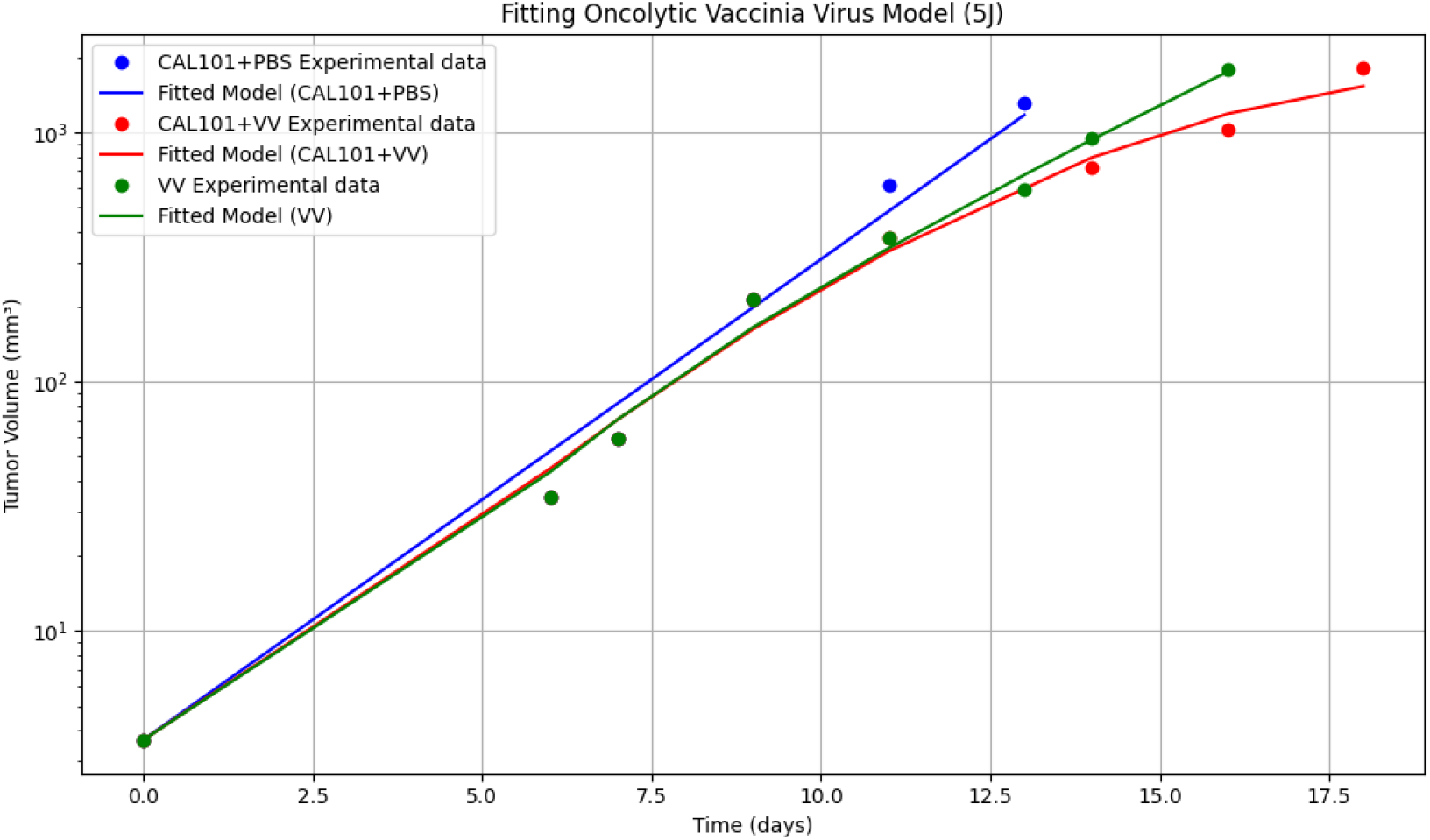
Model fits to experimental data from Figure 5J comparing CAL-101 treatment effects. Points show experimental data and lines show model fits for CAL-101+PBS (blue), VV alone (red), and CAL-101+VV combination treatment (green). The model captures the differential effects of CAL-101 on tumor progression.

Parameter estimates for the CAL-101 treatment groups (Table 3) revealed an intermediate phenotype between standard VV treatment and complete neutrophil depletion. Notably, the neutrophil clearance rate in CAL-101+VV treatment (1.88 *×* 10^−2^) fell between that of standard VV (2.22 *×* 10^−2^) and neutrophil-depleted conditions, suggesting partial inhibition of neutrophil function.

**Table 3:** Best fit parameter values and 95% confidence intervals for CAL-101 treatment conditions from Figure 5J analysis. Parameters are defined as in Table 2.

| Treatment | Parameter Values | 95% CI | SSR |
| --- | --- | --- | --- |
| CAL-101+PBS | $\lambda_T = 4.20 \times 10^{-1}$ | $1.94 \times 10^{-1} - 6.44 \times 10^{-1}$ | 0.0672 |
| | $\beta = 2.70 \times 10^{-4}$ | $1.00 \times 10^{-8} - 1.00 \times 10^{-2}$ | |
| | $\delta = 1.05 \times 10^{-4}$ | $8.32 \times 10^{-5} - 4.62 \times 10^{-1}$ | |
| | $p = 1.00 \times 10^1$ | $9.04 \times 10^0 - 1.19 \times 10^1$ | |
| | $c = 5.00 \times 10^{-2}$ | $2.26 \times 10^{-2} - 1.16 \times 10^{-1}$ | |
| | $k = 8.52 \times 10^{-4}$ | $1.00 \times 10^{-4} - 1.00 \times 10^0$ | |
| | $\gamma = 1.53 \times 10^{-1}$ | $1.00 \times 10^{-3} - 1.25 \times 10^0$ | |
| CAL-101+VV | $\lambda_T = 6.62 \times 10^{-1}$ | $2.68 \times 10^{-1} - 4.79 \times 10^0$ | 0.0463 |
| | $\beta = 3.37 \times 10^{-5}$ | $1.00 \times 10^{-8} - 1.00 \times 10^{-2}$ | |
| | $\delta = 1.00 \times 10^{-4}$ | $1.35 \times 10^{-5} - 1.85 \times 10^{-4}$ | |
| | $p = 1.00 \times 10^1$ | $2.23 \times 10^0 - 1.82 \times 10^1$ | |
| | $c = 5.00 \times 10^{-2}$ | $7.23 \times 10^{-3} - 8.70 \times 10^{-2}$ | |
| | $k = 1.88 \times 10^{-2}$ | $1.00 \times 10^{-4} - 1.00 \times 10^0$ | |
| | $\gamma = 8.20 \times 10^{-1}$ | $1.14 \times 10^{-2} - 6.01 \times 10^0$ | |

Bootstrap analysis of the CAL-101 treatment parameters (Figure 4) confirmed the intermediate effect on neutrophil function, showing parameter distributions between those of standard VV and neutrophil-depleted conditions.

**Figure 4:**
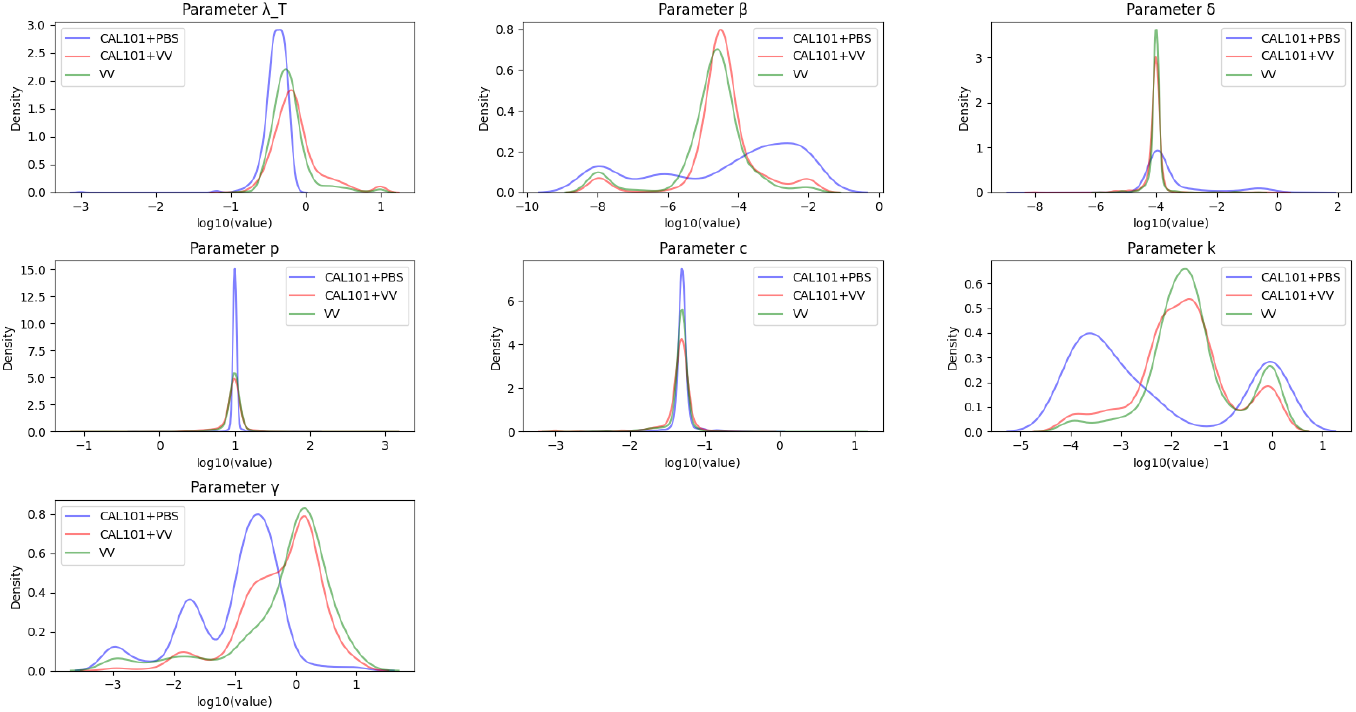
Distribution of bootstrapped parameter values across CAL-101 treatment conditions from Figure 5J analysis. Histograms show the logarithm of parameter values for CAL-101+PBS (blue), CAL-101+VV (red), and VV alone (green).

Parameter estimates for the CAL-101 treatment groups (Table 3) revealed an intermediate phenotype between standard VV treatment and complete neutrophil depletion. Notably, the neutrophil clearance rate in CAL-101+VV treatment (1.88 *×* 10^−2^) fell between that of standard VV (2.22*×* 10^−2^) and neutrophil-depleted conditions, suggesting partial inhibition of neutrophil function.

### 3.4 Statistical Analysis

Statistical comparison between treatment conditions using the Mann-Whitney U-test revealed significant differences in neutrophil-related parameters (Table 3.4). The neutrophil clearance rate (*k*) showed highly significant differences (*p* = 0.034, 78.0% of bootstrap iterations significant), while the tumor growth rate (*λ*_*T*_) showed no significant difference (*p* = 0.338), indicating that treatments primarily affect viral dynamics rather than intrinsic tumor growth.

**Table 4:**
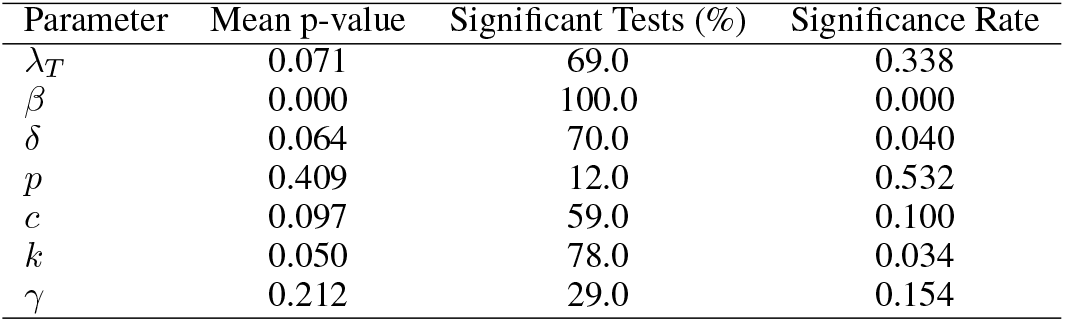
Statistical analysis of parameter differences between treatment conditions. Mean p-value represents the average from 1000 bootstrap iterations. Significant Tests shows the percentage of bootstrap iterations yielding p < 0.05.

| Parameter | Mean p-value | Significant Tests (%) | Significance Rate |
| --- | --- | --- | --- |
| $\lambda_T$ | 0.071 | 69.0 | 0.338 |
| $\beta$ | 0.000 | 100.0 | 0.000 |
| $\delta$ | 0.064 | 70.0 | 0.040 |
| $p$ | 0.409 | 12.0 | 0.532 |
| $c$ | 0.097 | 59.0 | 0.100 |
| $k$ | 0.050 | 78.0 | 0.034 |
| $\gamma$ | 0.212 | 29.0 | 0.154 |

## 4 Discussion

We find that an oncolytic vaccinia virus model without neutrophil-mediated clearance is unable to reproduce experimentally observed tumor progression patterns. The model parameterized only with basic virus-tumor interactions (*λ, β, δ*) fails to capture both the magnitude and timing of tumor response seen in the experimental data. The necessity of including neutrophil dynamics is supported by both our statistical analysis and other studies that have found immune components crucial for accurately modeling oncolytic virus therapy [Wodarz(2001), Komarova and Wodarz(2010), Friedman et al.(2006)].

The fitted parameters reveal striking differences between standard virus treatment (VV) and neutrophil-depleted conditions (LY6G+VV). Most notably, the viral clearance rate mediated by neutrophils (*k*) differs by four orders of magnitude between conditions (2.29 *×* 10^−6^ for VV vs. 1.45 *×*10^−10^ for LY6G+VV), highlighting the substantial impact of neutrophil depletion on viral persistence. This dramatic difference aligns with current understanding of neutrophil heterogeneity and their varied roles in viral clearance [Ng et al.(2019)]. The neutrophil response rate (*γ*) shows similarly dramatic variation (2.64 for VV vs. 2.63 *×* 10^−2^ for LY6G+VV), indicating successful modulation of neutrophil dynamics in the experimental system. These findings align with previous studies showing that immune-mediated viral clearance can be a major limiting factor in oncolytic virotherapy [Kim et al.(2015), Bajzer et al.(2008), Leliefeld et al.(2015)].

The comparison between different approaches to neutrophil modulation (LY6G antibody vs. CAL-101) provides additional insight into the mechanisms of treatment efficacy. While both approaches improve viral persistence compared to standard VV treatment, they show distinct parameter profiles. CAL-101 treatment results in intermediate values for neutrophil-related parameters, with viral clearance rates (5.54 *×* 10^−3^) falling between those of standard VV and LY6G+VV treatments. This hierarchical response is consistent with the known mechanisms of PI3K inhibition [Srinivasan et al.(2021), Yang et al.(2015)] and suggests that different mechanisms of neutrophil modulation may offer complementary therapeutic benefits, potentially informing future combination treatment strategies [Zhou et al.(2024)].

Bootstrap analysis of the parameter distributions reveals that the tumor growth rate (*λ*) remains relatively consistent across all treatment conditions, suggesting that neutrophil modulation primarily affects viral dynamics rather than intrinsic tumor properties. However, the viral production rate (*p*) shows significant variation (1.01 for VV vs. 1.00*×*10^−1^ for LY6G+VV), indicating that neutrophil activity may indirectly influence viral replication efficiency. The infection rate (*β*) also differs substantially between conditions (1.13 *×* 10^−7^ for VV vs. 3.70 *×* 10^−9^ for LY6G+VV), suggesting that neutrophil presence may affect viral spread within the tumor.

Statistical analysis of the model fits shows remarkably low SSR values across all experimental conditions (0.0350 for VV, 0.0853 for LY6G+VV, and comparable values for CAL-101 treatments), indicating good agreement between the model and experimental data. The slightly higher SSR values for neutrophil-depleted conditions might reflect increased biological variability in these modified systems, consistent with the complex role of neutrophils in regulating immune responses [Leliefeld et al.(2015)].

However, our model has several limitations that should be considered. While it captures neutrophil-mediated clearance, it does not account for other immune components such as natural killer cells or adaptive responses that may become more relevant in neutrophil-depleted conditions [Paiva et al.(2009)]. The model assumes spatial homogeneity, which may not fully represent the complex architecture of solid tumors and immune cell infiltration patterns [Morrow et al.(2022)]. Additionally, the different mechanisms of neutrophil modulation (antibody-mediated depletion vs. CAL-101 inhibition) may have distinct off-target effects that our model does not capture [Srinivasan et al.(2021)].

The timing of viral injections (days 8, 10, and 12) in our experimental protocol adds another layer of complexity to the system dynamics. Our model suggests that each injection leads to a temporary spike in viral concentration, but the rate of subsequent clearance differs markedly between treatment conditions. This observation has practical implications for optimizing treatment protocols, particularly regarding the timing and dosing of both viral administration and neutrophil modulation strategies.

These findings are particularly relevant given the current challenges in oncolytic virotherapy, where clinical response rates remain limited. Our results suggest that neutrophil modulation could significantly improve treatment outcomes, though careful consideration must be given to the timing and extent of immune modification. The dramatically lower *k* values in neutrophil-depleted conditions suggest that even partial neutrophil inhibition might substantially improve viral persistence and therapeutic efficacy. The complementary effects observed with different neutrophil modulation strategies further suggest that combination approaches might offer optimal therapeutic benefit.

Future studies should examine how these findings translate to human systems, as mouse models may not fully recapitulate human immune responses to viral therapy [Russell and Barber(2018)]. Investigation of potential synergies between different neutrophil modulation strategies, as well as combination with other immunomodulatory approaches, could provide valuable insights for improving oncolytic virotherapy outcomes [Morrow et al.(2022), Chen et al.(2021)]. Additionally, the development of more spatially-resolved models might better capture the complex dynamics of neutrophil infiltration and viral spread within the tumor microenvironment.

## Data Availability

The data presented in this study and all Python code used for analysis are available at: https://github.com/sahajsatani/oncolytic_vaccinia (accessed on 4 November 2024).

## Abbreviations The following abbreviations are used in this manuscript

VV: Vaccinia virus
ODE: Ordinary differential equation
SSR: Sum of squared residuals
CI: Confidence interval
LY6G: Lymphocyte antigen 6G
PBS: Phosphate buffered saline
FDA: Food and Drug Administration
T-VEC: Talimogene laherparepvec

## References

[Siegel et al.(2024)] Siegel, R.L.; Miller, K.D.; Jemal, A. Cancer Statistics, 2024. CA Cancer J. Clin. 2024, 74, 1–28.

[Cronin et al.(2023)] Cronin, K.A.; Lake, A.J.; Scott, S. Annual Report to the Nation on the Status of Cancer, Part I: National Cancer Statistics. Cancer 2023, 129, 837–884.

[Domingues et al.(2018)] Domingues, B.; Lopes, J.M.; Soares, P.; Populo, H. Melanoma Treatment in Review. Im-munoTargets Ther. 2018, 7, 35–49.

[MacKie et al.(2001)] MacKie, R.M.; Stewart, B.; Brown, S.M. Intralesional Injection of Herpes Simplex Virus 1716 in Metastatic Melanoma. Lancet 2001, 357, 525–526.

[Andtbacka et al.(2015)] Andtbacka, R.H.; Kaufman, H.L.; Collichio, F. Talimogene Laherparepvec Improves Durable Response Rate in Patients With Advanced Melanoma. J. Clin. Oncol. 2015, 33, 2780–2788.

[Ribas et al.(2017)] Ribas, A.; Dummer, R.; Puzanov, I. Oncolytic Virotherapy Promotes Intratumoral T Cell Infiltration and Improves Anti-PD-1 Immunotherapy. Cell 2017, 170, 1109–1119.

[Summers et al.(2010)] Summers, C.; Rankin, S.M.; Condliffe, A.M. Neutrophil Kinetics in Health and Disease. Trends Immunol. 2010, 31, 318–324.

[Drescher and Bai(2013)] Drescher, B.; Bai, F. Neutrophil in Viral Infections, Friend or Foe?Virus Res. 2013, 171, 1–7.

[Chen and Mellman(2017)] Chen, D.S.; Mellman, I. Elements of Cancer Immunity and the Cancer-Immune Set Point. Nature 2017, 541, 321–330.

[Wodarz(2001)] Wodarz, D. Viruses as Antitumor Weapons: Defining Conditions for Tumor Remission. Cancer Res. 2001, 61, 3501–3507.

[Komarova and Wodarz(2010)] Komarova, N.L.; Wodarz, D. ODE Models for Oncolytic Virus Dynamics. J. Theor. Biol. 2010, 263, 530–543.

[Friedman et al.(2006)] Friedman, A.; Tian, J.P.; Fulci, G.; Chiocca, E.A. Glioma Virotherapy: Effects of Innate Immune Suppression and Increased Viral Replication Capacity. Cancer Res. 2006, 66, 2314–2319.

[Kim et al.(2015)] Kim, P.S.; Crivelli, J.J.; Choi, I.K.; Yun, C.O.; Wares, J.R. Quantitative Impact of Immunomodulation versus Oncolysis with Cytokine-expressing Virus Therapeutics. Math. Biosci. Eng. 2015, 12, 841–858.

[Bajzer et al.(2008)] Bajzer, Ž.; Carr, T.; Josi’c, K.; Russell, S.J.; Dingli, D. Modeling of Cancer Virotherapy with Recombinant Measles Viruses. J. Theor. Biol. 2008, 252, 109–122.

[Zhou et al.(2024)] Zhou, H.; Li, W.; Chen, X.; Liu, L.; Yang, Z.; Wang, W.; Zhang, S.; Luo, Y.; Li, J.; Xie, C.; Hu, B.; Zhou, Y.; Guo, G.; Xie, X.; Wang, H. Neutrophils Limit Oncolytic Vaccinia Virus Therapy Through Lysosome-mediated Viral Degradation. Cancer Immunol. Immunother. 2024, 73, 453–467.

[Paiva et al.(2009)] Paiva, L.R.; Binny, C.; Ferreira, S.C.; Martins, M.L. A Multiscale Mathematical Model for Oncolytic Virotherapy. Cancer Res. 2009, 69, 1205–1211.

[Eftimie et al.(2011)] Eftimie, R.; Bramson, J.L.; Earn, D.J.D. Interactions Between the Immune System and Cancer: A Brief Review of Non-spatial Mathematical Models. Bull. Math. Biol. 2011, 73, 2–32.

[Ng et al.(2019)] Ng, L.G.; Ostuni, R.; Hidalgo, A. Heterogeneity of neutrophils. Nat. Rev. Immunol. 2019, 19, 255–265.

[Leliefeld et al.(2015)] Leliefeld, P.H.; Koenderman, L.; Pillay, J. How Neutrophils Shape Adaptive Immune Responses. Front. Immunol. 2015, 6, 471–482.

[Srinivasan et al.(2021)] Srinivasan, L.; Sasaki, Y.; Calado, D.P. PI3K Inhibition in B Cell Malignancies. Front. Immunol. 2021, 12, 701501.

[Yang et al.(2015)] Yang, Q.; Modi, P.; Newcomb, T.; Quelle, F.W. Idelalisib: First-in-Class PI3K Delta Inhibitor for the Treatment of Chronic Lymphocytic Leukemia, Small Lymphocytic Leukemia, and Follicular Lymphoma. Clinical Cancer Res. 2015, 21, 1537–1542.

[Morrow et al.(2022)] Morrow, R.J.; Brennan, N.A.; Scott, C.J. The Complex Role of the Tumor Microenvironment in Oncolytic Virus Therapy. Mol. Ther. 2022, 30, 2471–2483.

[Russell and Barber(2018)] Russell, S.J.; Barber, G.N. Oncolytic Viruses as Antigen-Agnostic Cancer Vaccines. Cancer Cell 2018, 33, 599–605.

[Chen et al.(2021)] Chen, P.; Liu, Z.; Liu, S. Current applications and future prospects of nanomaterials in tumor therapy. Int. J. Nanomedicine 2021, 16, 1937–1952.

